# Adolescent development of multiscale structural wiring and functional interactions in the human connectome

**DOI:** 10.1101/2021.08.16.456455

**Authors:** Bo-yong Park, Casey Paquola, Richard A. I. Bethlehem, Oualid Benkarim, Neuroscience in Psychiatry Network (NSPN) Consortium, Bratislav Mišić, Jonathan Smallwood, Edward T. Bullmore, Boris C. Bernhardt

## Abstract

Adolescence is a time of profound changes in the physical wiring and function of the brain. Here, we analyzed structural and functional brain network development in an accelerated longitudinal cohort spanning 14–25 years (n = 199). Core to our work was an advanced *in vivo* model of cortical wiring incorporating MRI features of cortico-cortical proximity, microstructural similarity, and white matter tractography. Longitudinal analyses assessing age-related changes in cortical wiring identified a continued differentiation of multiple cortico-cortical structural networks in youth. Studying resting-state functional MRI measures in the same participants at baseline, we found that regions with more similar structural wiring were more likely to be functionally coupled. Moreover, longitudinal structural wiring changes, particularly between sensory/unimodal and default mode networks, were reflected in tendencies for increased differentiation in brain function. These longitudinal findings provide new insights into adolescent development of human brain structure and function, illustrating how structural wiring interacts with the maturation of macroscale functional hierarchies.

## Introduction

In adolescence, increasing evidence suggests that ongoing maturation of structural and functional brain networks underpins broad cognitive development (1–9). Prior magnetic resonance imaging (MRI) literature has assessed regional changes in brain structure in youth (1–3, 10–16), showing age-related widespread decreases in cortical thickness (1, 13) as well as changes in surrogates of intracortical myelin content (15–17). Complementing these regional changes, diffusion MRI as well as functional MRI (fMRI) studies have shown an ongoing maturation of both the microstructure of inter-connecting white matter tracts as well as large-scale developmental changes in functional organization, indicative of shifts in brain connectivity towards a more distributed network topology (18–20). Utilizing multimodal longitudinal MRI analyses, here, we explored how adolescent structural network development gives rise to potential shifts in functional network architecture.

Core to our work was a comprehensive and recently introduced *in vivo* model of cortical wiring, which integrates several neuroimaging features of structural connectivity *i.e*., diffusion MRI tractography, cortico-cortical geodesic distance mapping, and microstructural covariance analysis (21). Diffusion MRI tractography maps white matter fibers, showing increasing validity in approximating deeper tracts, but some limitations in the proximity of cortical grey matter regions (22, 23). On the other hand, geodesic distance analysis measures the spatial proximity of areas across the cortical sheet, tapping into short range cortico-cortical connectivity and wiring cost (24). Finally, a recent extension of structural covariance analysis (25, 26), labelled microstructural profile covariance analysis, identifies networks with similar myelin-sensitive imaging characteristics across cortical depths in a subject-specific manner (27, 28). By integrating these complementary measures from diffusion MRI tractography, geodesic distance, and microstructural covariance via unsupervised pattern learning, we could generate a new coordinate system and arrange cortical regions with respect to their similarity in structural wiring (21). In a prior evaluation in healthy adults, we demonstrated that this novel approach captures spatial gradients of (i) cortical cytoarchitecture, (ii) cell-type specific gene expression, and (iii) intrinsic functional connectivity and signal flow measured from resting-state fMRI (rs-fMRI) and intracranial electrical recordings (21), supporting neurobiological and functional validity. Here, we adopted this wiring model to chart adolescent development of cortical structural networks longitudinally.

As brain structure ultimately scaffolds brain function (29–35), it is not surprising that multiple functional networks also change throughout adolescence. Prior analyses based on rs-fMRI connectivity analysis in youth have shown marked shifts in the connectivity patterns of multiple large-scale cortical networks. A particular emphasis has been placed on the default mode and frontoparietal networks, both known to be spatially distributed and important for higher cognition (3, 36, 37). In one recent study, it was furthermore shown that different cortical areas undergo variable functional maturational trajectories, differentiating sensory and motor networks that follow a more ‘conservative’ functional trajectory from transmodal systems such as the default mode and frontoparietal networks that show a more ‘disruptive’ mode, reflected by reconfiguration of their functional connectivity patterns towards a more distributed and long-range network architecture (37). Benefitting from an increasing availability of multimodal datasets, several studies have begun to examine how brain structure and function co-mature. For example, a prior study showed that structural network modules become more segregated with advancing age, and that this process reflects ongoing development of executive function from 8 to 22 years (38). In other studies, the authors showed changes in structurefunction coupling with ongoing age, particularly with respect to transmodal vs sensory and motor networks (3, 39). The current work built on this growing literature to assess how adolescent changes in cortical wiring are reflected in functional network maturation.

Our study was based on the Neuroscience in Psychiatry Network (NSPN) 2400 cohort, an accelerated longitudinal dataset that enrolled healthy individuals between 14–25 years (16, 40). Structural wiring models were derived for each participant at two time points based on multimodal neuroimaging and unsupervised machine learning (41), and we estimated longitudinal trajectories in structural network maturation using linear mixed effect models. In addition to assessing whether age-effects on structural wiring were similar to cortical thickness changes in the same subjects (1, 11, 13, 42), we examined how structural wiring changes reflect adolescent functional network maturation based on parallel rs-fMRI acquisitions. Multiple sensitivity analyses assessed robustness of our findings with respect to several analysis parameter variations.

## Results

We studied 199 healthy participants obtained from the NSPN 2400 cohort, who were part of the accelerated longitudinal design and had imaging data available (16, 40) (**Fig. 1A**). Included participants had two measurement time points (mean inter-scan interval was 0.94 years, range = 0.5–1), with a mean age of 18.84 (range = 14–25) years at baseline and 19.96 (range = 15–26) years at follow-up. Participants were uniformly distributed across the entire age range, with a similar sex ratio (52/48% males/females). Participant demographics, image processing, and analysis are further detailed in the *Methods*.

**Fig. 1.**
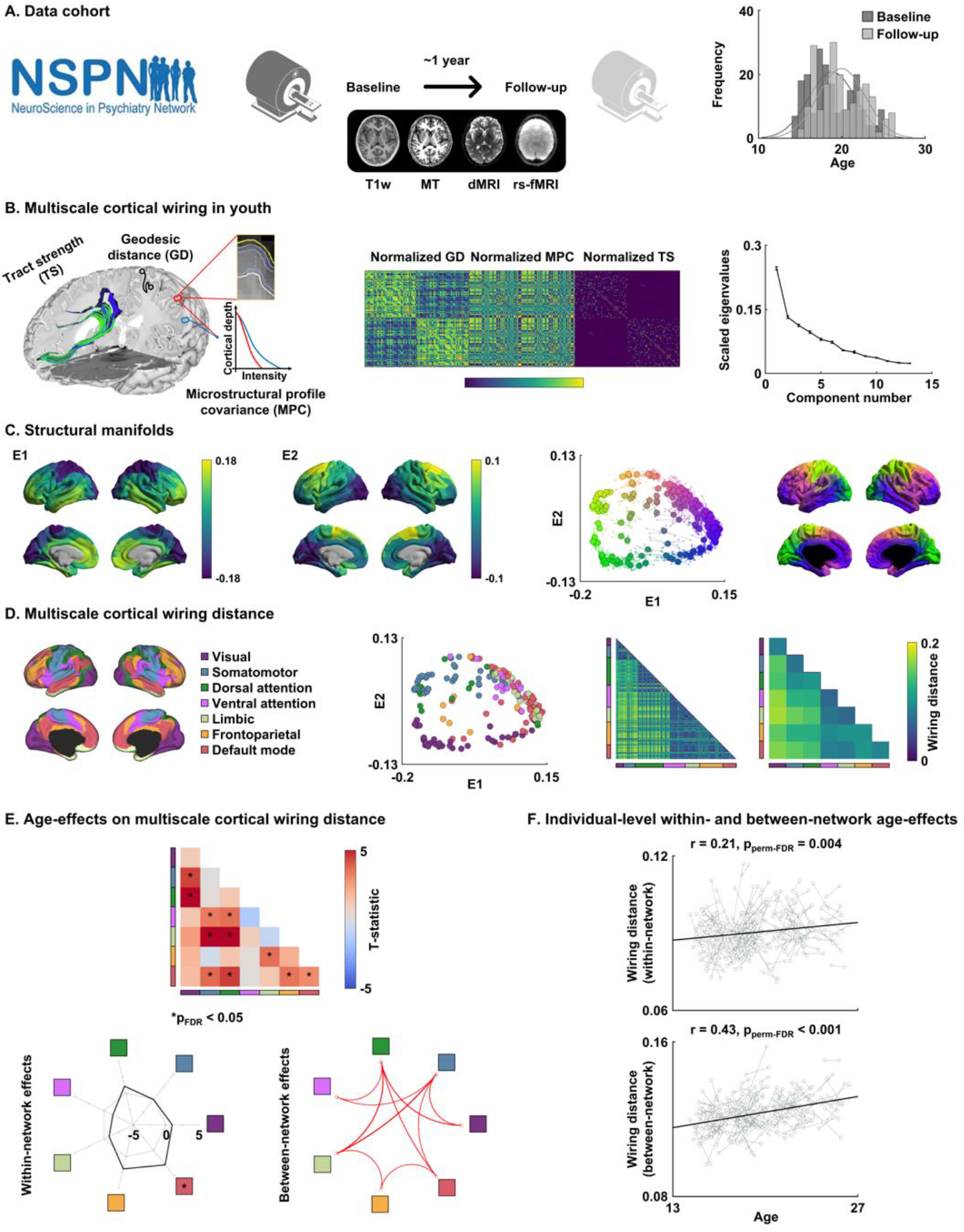
Adolescent development of multiscale cortical wiring. **(A)** We studied the multimodal MRI dataset from the NSPN 2400 cohort, examining both baseline and follow-up scans. Age at both visits is represented in the histograms. **(B)** Our *in vivo* model of cortico-cortical wiring combined three cortical neuroimaging features *i.e*., geodesic distance (GD), microstructural profile covariance (MPC), and tract strength (TS). Matrices were normalized and concatenated prior to applying non-linear manifold learning, which identifies a coordinate system informed by cortical wiring. The scree plot shows eigenvalues of each estimated component, with error bars indicating the SD across ten repetitions. **(C)** We estimated two eigenvectors (E1, E2) from cortical wiring features. Averaged maps across ten repetitions are reported. The scatter plot represents each brain region projected onto the two-dimensional wiring space with different colors, mapped onto the cortical surface. Solid dots indicate mean across ten repetitions, and transparent dots with lines indicate results from each repetition. **(D)** Nodes in the wiring space were assigned to seven intrinsic functional communities. Multiscale cortical wiring distance, *i.e*., the Euclidean distance between different nodes in the wiring space, was calculated at a node-level and summarized for intrinsic functional communities. **(E)** The t-statistics of age-effects on cortical wiring distance within- and between-networks are reported, with significant (p_FDR_ < 0.05) results marked with asterisks. The within-network effects are represented with radar plots, and significant networks are reported with asterisks. Significant between-network effects are reported with circular plots. **(F)** The scatter plots show age-related changes in within (upper) and between-network (lower) wiring distance of each individual in the identified networks. *Abbreviation:* NSPN, the Neuroscience in Psychiatry Network; T1w, T1-weighted; MT, magnetization transfer; dMRI, diffusion magnetic resonance imaging; rs-fMRI, resting-state functional magnetic resonance imaging; FDR, false discovery rate.

### Multiscale cortical wiring in youth

Following a recently developed approach in healthy adults (21), we built a comprehensive *in vivo* model of cortico-cortical wiring for every subject time point (**Fig. 1B**). Models combined MRI-based measures of geodesic distance (GD), microstructural profile covariance (MPC), and diffusion MRI tract strength (TS). We integrated these three complementary features into a common lowdimensional space using a non-linear manifold learning technique (see *Methods*) (41). Two eigenvectors (E1, E2) were identified that collectively explained approximately 37.8 ± 0.01% (mean ± SD) of information, and averaged across ten iterations with different non-overlapping subsets within the NSPN cohort (**Fig. 1B–C**; see *Methods*). The first eigenvector (E1) depicted a sensory-fugal gradient from sensory/motor towards transmodal networks such as the default mode and frontoparietal networks and the second eigenvector (E2) differentiated anterior and posterior cortices. We calculated the Euclidean distance between all brain regions in the wiring-derived low dimensional space, as a measure of structural differentiation (henceforth *wiring distance*; **Fig. 1D**; see *Methods* for details). While within-network connectivity showed overall low wiring distance, connections between sensory and transmodal regions showed high values. In other words, the wiring distance measure captured an overall integration of nodes involved in the same functional network and an overall segregation between networks, particularly between sensory *vs* transmodal networks. Findings were furthermore summarized according to intrinsic functional communities (43).

### Tracking adolescent changes in multiscale cortical wiring

We assessed age-effects on this wiring distance using linear mixed effect models. In adolescence, several prior studies have reported robust age-related changes in cortical thickness (1, 11, 13, 42), and we confirmed similar age-effects in our NSPN cohort. Indeed, cortical thickness decreased in widespread cortical regions with advancing age (false discovery rate (p_FDR_) < 0.05; **Fig. S1A**). Running a spatial correlation analysis between longitudinal thickness and wiring effects, while controlling for spatial autocorrelation of the two maps (44), we only observed a weak spatial association to changes in wiring measures, with no correlation to within-network wiring distance (r = −0.02 ± 0.05, p_spin-FDR_ = 0.27) and a trend-level association to between-network wiring distance (r = −0.12 ± 0.04 across ten repetitions, p_spin-FDR_ = 0.06; **Fig. S1B**). We then assessed age-effects on wiring distance after controlling for sex, site, head motion, and subject-specific random intercepts, as well as cortical thickness (45). We found robust increases in wiring distance within and across multiple networks with advancing age (p_FDR_ < 0.05; **Fig. 1E**), indicative of an ongoing structural differentiation in multiple systems in youth. Among the seven large-scale communities, the default mode network showed significant within-network changes, and we furthermore observed increased wiring distance across several between-network connections, particularly between nodes of default mode and attention, frontoparietal regions as well as between sensory and attention, limbic networks (p_FDR_ < 0.05). Investigating changes in wiring distance for each individual in the identified networks, we could show a low but significant association between age and within-network wiring distance (r = 0.21, p_perm-FDR_ = 0.004) as well as a moderate association between age and between-network wiring distance (r 0.43, p_perm-FDR_ < 0.001; **Fig. 1F**).

We additionally assessed age-effects on each cortical feature (*i.e*., GD, MPC, and TS) to quantify how wiring distance captures age-related changes in cortical organization relative to changes in single features (**Fig. S2**). When analyzing wiring distance, the effect size (*i.e*., the mean absolute t-statistic across network pairs) was 32.07 ± 17.88% higher than when studying only GD across ten repetitions (see *Methods*), 15.45 ± 6.53% higher than when studying MPC, and 14.65 ± 11.58% higher than when studying TS, indicating that wiring distance describes adolescent cortical reorganization more sensitively than each modality separately. When associating age-effects on wiring distance with those on each feature, wiring distance increases were strongly related to reductions in MPC (r = −0.58, p_FDR_ = 0.001), but not very much to changes in TS (r = −0.25, p_FDR_ = 0.21) nor GD (r = 0.22, p_FDR_ = 0.27), supporting the notion that increases in multiscale wiring distance reflected mostly a decreased similarity of intracortical microstructure. Furthermore, as the overall manifold size (*i.e*., mean of wiring distance across all networks) was significantly associated with age (r = 0.19 ± 0.02, p < 0.001 across ten repetitions), we repeated linear mixed effect models after additionally controlling for mean wiring distance (**Fig. S3**). Despite decreases in the effect size, we observed overall consistent patterns, confirming that age-effects on wiring distance were not driven by the expansion in manifold space itself.

### Associations with macroscale functional network maturation

To evaluate functional associations of the changes in multiscale cortical wiring, we first generated functional connectivity based on rs-fMRI obtained in the same subjects at equivalent time points (**Fig. 2A**). Associating structural cortical wiring distance and functional connectivity across intrinsic functional networks at a cross-sectional level, we found strong negative structure-function coupling (r = −0.74, p_spin_ < 0.001; **Fig. 2B**). In other words, regions with increased wiring distance generally show weaker functional connectivity. We then charted the development of functional connectivity across age, and we found decreases in sensory-default mode network connectivity and increases in connectivity between sensory networks (p_FDR_ < 0.05; **Fig. 2C**). To assess how the age-related changes in structural and functional measures were inter-related, we correlated the age-effects on wiring distance with the age-effects on functional connectivity. Here, we found a tendency for a negative association (r = −0.21, p_spin_ = 0.09; **Fig. 2D**). The results indicate that cross-sectionally, weaker interconnectivity in brain function between sensory and transmodal networks is coupled with higher cortical wiring distance between these networks. Moreover, age-related functional differentiation between sensory and default mode networks also tends to be reflected in increased differentiation in structural wiring during adolescence.

**Fig. 2.**
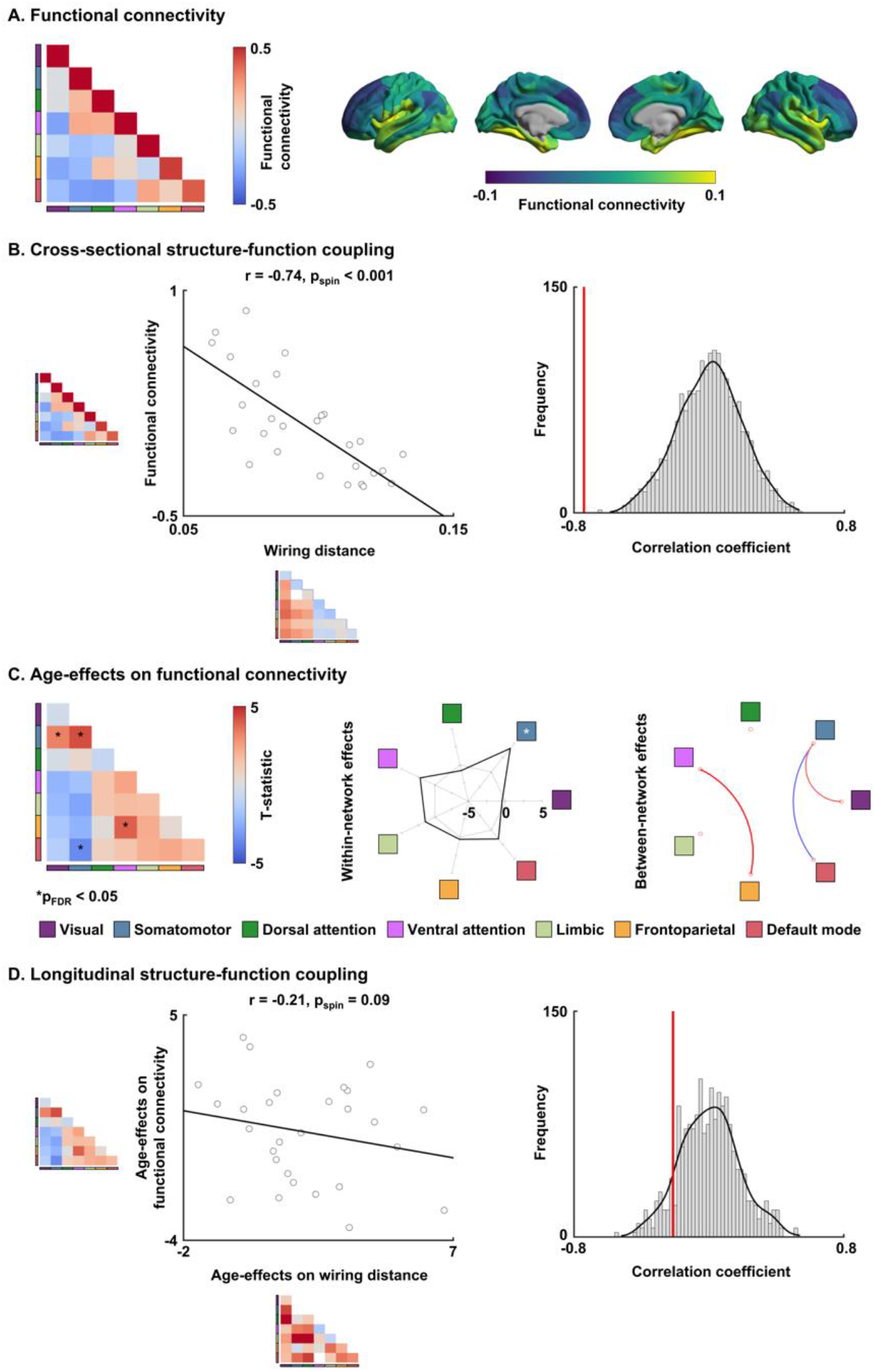
Association between functional connectivity and wiring distance. **(A)** Functional connectivity matrix was summarized according to intrinsic functional communities (left) and projected onto brain surfaces (right). **(B)** Cross-sectional structure-function coupling between functional connectivity and cortical wiring distance. The histogram indicates distribution of correlation coefficients, and the actual r-value is represented with a red bar. **(C)** Age-effects on functional connectivity. The t-statistics of age-effects are reported. The within-network effects are represented with radar plots, and significant networks are reported with asterisks. Significant between-network effects are reported with circular plots. **(D)** Longitudinal structure-function coupling between age-effects on functional connectivity and cortical wiring distance. *Abbreviation:* FDR, false discovery rate.

### Sensitivity analysis

We assessed whether our findings were robust with respect to several methodological variations.

#### a) Parcellation scales

We repeated assessing age-effects using different parcellation scales (*i.e*., 100 and 300 regions) and revealed consistent results (**Fig. S4**), indicating robustness of our findings across different scales.

#### b) Structural manifold generation using principal component analysis

Our main analysis estimated structural manifolds using diffusion map embedding (46), in keeping with a previous approach to study structural manifolds in healthy young adults (21, 47). We repeated our analysis after alternatively estimating structural manifolds using principal component analysis (48), and the manifolds and age-effects were similar (**Fig. S5**), confirming robustness.

#### c) Parcellation scheme

We generated connectome manifolds and assessed adolescent remodeling using a functional (*i.e*., Schaefer) parcellation (49) instead of structural parcellation scheme (50), and found consistent results (**Fig. S6**), indicating the robustness of our analyses across different parcellations.

## Discussion

The current work assessed adolescent maturation of cortical networks based on an advanced *in vivo* model of cortical wiring (21). Charting typical development from late childhood to early adulthood using the longitudinal NSPN cohort (16, 40), we observed marked increases in within- and between-network wiring distances in both sensory and transmodal association networks, indicating of ongoing structural differentiation in youth across multiple cortical networks. Moreover, associating cortical structural wiring features with intrinsic functional connectivity obtained from parallel rs-fMRI analysis performed in the same subjects, we observed that functional networks reconfigure alongside the marked reorganization of cortico-cortical wiring. Collectively, our work offers a novel perspective on how structural brain networks reconfigure and how these changes give rise to ongoing functional maturation in typically developing youth.

Our work centered on an advanced *in vivo* model of structural wiring that integrates multiple dimensions of cortico-cortical connectivity (21), namely diffusion MRI tractography strength (TS), geodesic distance (GD), and microstructure profile covariance (MPC). Each feature taps into different aspects of cortico-cortical connectivity, grounded in seminal neuroanatomical work on the multiple facets of the cortical wiring scheme (51). Synergistic integration of these features is hypothesized to comprehensively describe structural connectivity, and to thus reveal structure-function relationships in the developing brain. In fact, TS is an established measure of short- and long-range fibers in the white matter (52, 53), whereas GD is computed within the cortical ribbon, approximating horizontal connectivity between adjacent cortical regions (24, 54). Similarity of intracortical microstructural profiles, quantified as MPC (28), is also recognized as an indicator of inter-regional connectivity (21, 27, 55, 56). In fact, the structural model of brain connectivity, initially formulated in non-human animals, predicts that areas with similar microstructure are more likely to be connected than areas with different connectivity profiles (57). These findings were recently extended to human neuroanatomy, by relating microstructural similarity to diffusion MRI-derived streamline strength (56, 58) and to resting-state functional connectivity (28, 59). Here, we fused and mapped the three above cortical wiring features into a 2D coordinate system using manifold learning techniques (21, 41, 46). By translating the approach previously formulated in adults (21) to typically developing adolescents, we demonstrated that the wiring space in youth overall resembles the one previously seen in adults. Indeed, the two principal dimensions of the wiring space represented sensory-fugal and anterior-posterior gradients – two major axes of adult macroscale cortical topography (60–64). On the other hand, we could obtain new insights into adolescent reconfigurations of structural networks via longitudinal analyses. The NSPN dataset was built using an accelerated longitudinal design, enrolling individuals aged from late childhood to young adulthood with a 1 year follow up on average (16, 40). Compared to cross-sectional studies, longitudinal designs measure intra-individual changes in cortical features and chart developmental trajectories directly (3, 20, 65–67). Our multiscale approach gathered evidence for developmental shifts in cortical wiring, indicative of increased wiring distances in multiple systems of the cortical mantle, with highest effects in default mode and ventral attention networks. These findings indicate a continued differentiation of cortico-cortical structural networks, which most markedly take place in transmodal systems at the apex of the cortical hierarchy (3, 37, 68, 69). Notably, wiring space analysis revealed increased effects compared to analysis of single features, suggesting that our compact multiscale approach may offer additional sensitivity in the study of adolescent development. These findings could, thus, recapitulate prior work in adolescence more generally and the NSPN dataset specifically, including our recent work showing overall changes in cortical myelination (16, 70) as well as depth-dependent shifts in intracortical myeloarchitecture (17). Moreover, several studies have described structural connectivity changes based on diffusion MRI tractography, reporting general increases in streamline strength in transmodal areas in adolescence (20, 71), together with enhanced within-module integration and strengthening of structural network hubs, sometimes alongside a weakening of more local connections (72, 73). In our work, different constituent wiring features contributed in a graded manner to our overall findings, with a marked association wiring distance increases and ongoing microstructural differentiation of transmodal areas from the rest of the brain (17).

Alterations in cortical morphology during adolescence are well established, and the prevailing findings in the literature indicate widespread cortical thickness reductions with advancing age, a finding likely reflecting ongoing synaptic pruning and cortical myelination (10, 16, 74, 75). Here, by analyzing longitudinal cortical thickness changes in the same NSPN participants, we could confirm widespread cortical thinning in youth with advancing age. What’s more, we showed that wiring space changes were only partially attributable to these changes in cortical thickness, suggesting that age-related structural wiring changes likely occurred above and beyond maturational effects on cortical morphology per se. In prior work in healthy adults (21), we could identify associations between *in vivo* cortical wiring space organization and intracortical factors, specifically cell-type specific gene expression as well as externopyramidization. Although these associations were indirect and based on separate datasets (*in vivo* MRI and histology-based *post mortem* gene expression information), they nevertheless supported a link between multiscale wiring and internal cortical microcircuitry that go beyond the changes measurable by cortical thickness measures alone. Such interactions between different scales of cortical organization during typical development could be further explored in studies obtaining wiring space data and gene expression in the same subjects.

During adolescence, the age-related reconfiguration in functional connectome organization has recently been shown to mainly follow two distinct trajectories, labeled as *conservative* and *disruptive* modes (37). Conservative modes involve the ongoing strengthening of already strong functional connectivity and primarily take place in primary cortical regions. On the other hand, disruptive functional maturational trajectories have been observed in cortical association areas and subcortical nodes, and are characterized by a strengthening of initially weak connections or as a weakening of initially strong connections (37). These results complement our structural wiring space findings of an alteration of functional network topologies in adolescence, which showed increases in wiring distance between sensory and transmodal regions. Furthermore, assessing spatial associations between age-effects on structural wiring and functional connectivity, we observed that adolescent decreases in functional connectivity between sensory and association systems are marginally reflected in increased cortical wiring distances between these systems. These segregation patterns of sensory-transmodal networks echo prior studies in individuals aged 8-23 that has shown marked reconfigurations of structure-function coupling during development, particularly in sensory and transmodal regions, which further could support increased functional flexibility and cognitive control with advancing age during that time window (3, 38, 76).

To conclude, we tracked longitudinal development of structural brain networks in adolescence based on an advanced model of cortical structural wiring, and showed an ongoing differentiation in cortico-cortical wiring across multiple brain networks. Parallel analysis of rs-fMRI connectivity data obtained in the same subjects could show ongoing maturation of functional networks, which tended to reflect the observed changes in structural wiring. Our multimodal framework, thus, provides novel insights into structural and functional brain development in adolescence, and points to an inherent coupling of developmental trajectories across both domains.

## Methods

### Participants

We obtained imaging and phenotypic data from the NSPN 2400 cohort, which contains questionnaire data on 2,402 individuals (with MRI data in a subset of ~300) from adolescence to young adulthood in a longitudinal setting (16, 40). In this study, we included 199 participants who completed quality-controlled (see *Data preprocessing* section) multimodal MRI scans consisting of T1-weighted, magnetization transfer (MT), diffusion MRI, and rs-fMRI for at least two time points (48% female; mean ± SD age = 18.84 ± 2.83 (between 14 and 25) years at baseline and 19.96 ± 2.84 (between 15 and 26) years at follow-up with inter-scan interval of 0.94 ± 0.17 (between 0.5 and 1) years; **Fig. 1A**). Data were collected from three different sites: Wolfson Brain Imaging Centre; MRC Cognition and Brain Sciences Unit in Cambridge; and University College London. Participants provided informed written consent for each aspect of the study, and parental consent was obtained for those aged 14–15 years old. Ethical approval was granted for this study by the NHS NRES Committee East of England-Cambridge Central (project ID 97546). The authors assert that all procedures contributing to this work comply with the ethical standards of the relevant national and institutional committees on human experimentation and with the Helsinki Declaration of 1975, as revised in 2008.

### MRI acquisition

Imaging data were obtained using a Siemens Magnetom TIM Trio 3T scanner at all sites. The T1-weighted and MT sequences were acquired using a quantitative multiparameter mapping (MPM) sequence (repetition time (TR)/flip angle = 18.7ms/20° for T1-weighted and 23.7ms/6° for MT; six equidistance echo times (TE) = 2.2–14.7ms; voxel size = 1mm^3^; 176 slices; field of view (FOV) = 256 × 240mm; matrix size = 256 × 240 × 176) (77). The diffusion MRI data were acquired using a spin-echo echo-planar imaging (EPI) sequence (TR = 8,700ms; TE = 90ms; flip angle = 90°; voxel size = 2mm^3^; 70 slices; FOV = 192 × 192mm^2^; matrix size = 96 × 96 × 70; b-value = 1,000s/mm^2^; 63 diffusion directions; and 6 b0 images). The rs-fMRI data were collected using a multi-echo EPI sequence with three different TEs (TR = 2.43 ms; TE = 13.0/30.55/48.1 ms; flip angle = 90°; voxel size = 3.75 × 3.75 × 4.18 mm^3^; 34 slices; FOV = 240 × 240 mm^2^; matrix size = 64 × 64 × 34; and 269 volumes).

### Data preprocessing

T1-weighted data were processed using the fusion of neuroimaging preprocessing (FuNP) pipeline integrating AFNI, FSL, FreeSurfer, ANTs, and Workbench (https://gitlab.com/by9433/funp) (78–82), which is similar to the minimal preprocessing pipeline for the Human Connectome Project (83). Gradient nonlinearity and b0 distortion correction, non-brain tissue removal, and intensity normalization were performed. The white and pial surfaces were generated by following the boundaries between different tissues (80, 84–86). The midthickness surface was generated by averaging the white and pial surfaces, and it was used to generate an inflated surface. Quality control involved visual inspection of surface reconstruction of T1-weighted data, and cases with faulty cortical segmentation were excluded. Surface-based co-registration between T1-weighted and MT weighted scans were performed. We generated 14 equivolumetric cortical surfaces within the cortex, especially between inner white and outer pial surfaces, and sampled MT intensity along these surfaces (28). The diffusion MRI data were processed using MRtrix3 (23), including correction for susceptibility distortions, head motion, and eddy currents. The rs-fMRI data were processed using multi-echo independent component analysis (ME-ICA) pipeline (https://github.com/ME-ICA/me-ica) (87, 88). The first six volumes were discarded to allow for the magnetic field saturation, and slice timing was corrected. Motion correction parameters were estimated from the middle TE data by aligning all volumes to the first volume using rigid-body transformation. The co-registration transformation parameters from functional to structural image were estimated by registering the skullstripped spatially concatenated multi-echo functional data to the skull-stripped anatomical image using affine transformation. The estimated motion correction and anatomical co-registration parameters were applied to each slice-timing corrected TE data and then temporally concatenated. The noise components were removed using principal component analysis followed by independent component analysis (87, 88). The processed fMRI data were mapped to the standard grayordinate space (*i.e*., 32k Conte69) with a cortical ribbon-constrained volume-to-surface mapping algorithm. Finally, data were surface smoothed with 5 mm full width at half maximum.

### Multiscale cortical wiring features

We calculated complementary cortical wiring features from different imaging sequences, namely GD from T1-weighted, MPC from MT, and TS from diffusion MRI (**Fig. 1B**). GD is a physical distance represented by the shortest paths between two points along the cortical surface (24, 47, 54). To calculate the GD matrix, we first matched each vertex to the nearest voxel in volume space. Then we calculated the distance to all other voxels traveling through a grey/white matter mask using a Chamfer propagation (https://github.com/mattools/matImage/wiki/imGeodesics) (89). Unlike a previously introduced approach that calculates only intra-hemispheric distance (24, 47, 54), this approach allows estimating interhemispheric projections (21). We mapped GD to 200 cortical nodes parcellation scheme, which preserves the boundaries of the Desikan Killiany atlas (50). Following our prior study in adults (28), the MPC matrix was constructed by calculating linear correlation of cortical depth-dependent intensity profiles between different nodes, controlling for the average whole-cortex intensity profile based on the 200 parcels. The MPC matrix was thresholded at zero and log-transformed. We generated the TS matrix from preprocessed diffusion MRI data using MRtrix3 (23). Anatomical constrained tractography was performed using different tissue types derived from the T1-weighted image, including cortical and subcortical grey matter, white matter, and cerebrospinal fluid (90). We estimated co-registration transformation from T1-weighted to diffusion MRI data with boundary-based registration and applied the transformation to different tissue types to align them onto the native diffusion MRI space. The multi-shell and multi-tissue response functions were estimated (91), and constrained spherical deconvolution and intensity normalization were performed (92). Seeding from all white matter voxels, the tractogram was generated using a probabilistic approach (23, 93) with 40 million streamlines, a maximum tract length of 250, and a fractional anisotropy cutoff of 0.06. Subsequently, we applied spherical-deconvolution informed filtering of tractograms (SIFT2) to optimize an appropriate cross-section multiplier for each streamline (94), and reconstructed wholebrain streamlines weighted by cross-section multipliers. Reconstructed cross-section streamlines were mapped onto the 200 parcels to build TS matrix, and log-transformed (95, 96).

### Structural manifold identification

We estimated structural manifolds based on the multiscale cortical features calculated above using an openly accessible normative manifold map approach (https://github.com/MICA-MNI/micaopen/tree/master/structural_manifold) (21), which is now integrated in BrainSpace (https://github.com/MICA-MNI/BrainSpace) (41). First, we rank normalized nonzero entries of the input matrices, and the less sparse matrices (*i.e*., GD and MPC) were rescaled to the same numerical range as the sparsest matrix (*i.e*., TS) to balance the contribution of each input measure (**Fig. 1B**). Notably, we rank normalized the inverted GD matrix to represent closer regions with larger values. We horizontally concatenated the normalized GD, MPC, and TS matrices and constructed an affinity matrix with a normalized angle kernel with 10% density, which quantifies the strength of cortical wiring between two regions. Structural manifolds were estimated via diffusion map embedding (46) (**Fig. 1C**), which is robust to noise and computationally efficient compared to other non-linear manifold learning techniques (97, 98). It is controlled by two parameters α and *t*, where α controls the influence of the density of sampling points on the manifold (α = 0, maximal influence; α = 1, no influence) and *t* controls the scale of eigenvalues of the diffusion operator. We set α = 0.5 and t = 0 to retain the global relations between data points in the embedded space, following prior applications (17, 20, 28, 41, 47, 99, 100). Cortical regions with more similar inter-regional patterns are more proximal in this new structural manifold. To assess robustness, we repeated estimating structural manifolds ten times with different sets of participants. Specifically, we split the dataset into nonoverlapping template (1/10) and non-template (9/10) partitions with similar distribution of age, sex, and site. The template manifold was generated using the averaged concatenated matrix of template dataset, and individual-level manifolds were estimated from the non-template dataset and aligned to the template manifold via Procrustes alignment (41, 101). We repeated generating connectome manifolds ten times with different template and non-template datasets.

### Age-effects on structural manifolds

To chart age-effects on structural manifolds, we first calculated multiscale cortical wiring distance, which is the Euclidean distance between different brain regions in the manifold space (**Fig. 1D**) (21, 102), and stratified the node-level wiring distance based on intrinsic functional communities (43). It has been shown that cortical thickness shows significant changes across age (1, 11, 13, 42). We first replicated these morphological findings by assessing age-effects on cortical thickness measured using T1-weighted MRI (**Fig. S1A**). Next, we linearly correlated time-related changes in wiring distance and those in cortical thickness to assess spatial similarity across the cortex (**Fig. S1B**). The significance of the similarity was assessed based on 1,000 spin tests that account for spatial autocorrelation (41, 44), and FDR corrected across within and between-network correlations. We then assessed age-effects on network-level wiring distance using a linear mixed effect model (45). The model additionally controlled for sex, site, head motion (*i.e*., frame-wise displacement measured from diffusion MRI), cortical thickness, and included a subject-specific random intercept. We corrected for multiple comparisons across all pairs of functional communities with p_FDR_ < 0.05 (103). We repeated the age modeling ten times with different non-template individuals and reported only those network pairs showing significant effects across all repetitions (**Fig. 1E**). To assess individual-level changes in wiring distance across the age, we calculated linear correlations between mean age and within- and between-network wiring distance in the identified networks between baseline and follow-up, where the significance was determined based on 1,000 permutation tests randomly assigning subjects (**Fig. 1F**). We additionally implemented mixed effect models for each cortical wiring feature separately (*i.e*., GD, MPC, and TS) to assess how much the age-effects improved when we considered multiscale cortical wiring distance (**Fig. S2**). The age-effect t-statistics of each feature were correlated with those of wiring distance to assess which features are strongly related to adolescent development in wiring distance. To assess the association between global manifold effects and age, we calculated linear correlation between age and mean wiring distance across the whole network. We also implemented a linear mixed effect model that additionally controlled for mean wiring distance to assess whether the age-effects on wiring distance are affected by global changes in the size of manifold space (**Fig. S3**).

### Association between structural manifolds and functional connectivity

Structure-function coupling analyses assessed how multiscale cortical wiring related to functional connectivity. First, we constructed the functional connectivity matrix by calculating linear correlations of resting-state functional time series between different brain regions, controlling for average whole-cortex signals (**Fig. 2A**). After row-wise thresholding remaining 10% of values for each row in the connectivity matrix, we assessed structure-function correspondence by computing linear correlations between the z-transformed functional connectivity and wiring distance at networklevel (43) (**Fig. 2B**). To assess the relationship between age-effects on multiscale cortical wiring and those on functional connectivity, we first assessed age-effects on functional connectivity to obtain node by node t-statistics (**Fig. 2B**). Then, we calculated linear correlation between the age-effect t-statistics of functional connectivity and wiring distance (**Fig. 2D**). We calculated correlations 1,000 times with spin test (41, 44).

### Sensitivity analysis

#### a) Parcellation scales

Our main analyses were based on the structural atlas of 200 cortical nodes defined using Desikan Killiany atlas (50). To assess robustness across multiple parcellation scales, we generated structural manifolds using structural atlases with 100 and 300 parcels and repeated the age modeling (**Fig. S4**).

#### b) Structural manifold generation using principal component analysis

Instead of relying on diffusion map embedding (46), we generated structural manifolds using principal component analysis (48). Then, we repeated calculating multiscale cortical wiring distance and assessed age-effects to evaluate consistency of our findings (**Fig. S5**).

#### c) Functional parcellation

We also repeated structural manifold generation and age modeling using the functional Schaefer parcellation scheme with 200 parcels (49) (**Fig. S6**).

## Data and code availability

The imaging and phenotypic data were provided by the Neuroscience in Psychiatry Network (NSPN) 2400 cohort. As stated in https://doi.org/10.1093/ije/dyx117, the NSPN project is committed to make the anonymised dataset fully available to the research community, and participants have consented to their de-identified data being made available to other researchers. A data request can be made to. Codes for multimodal connectome manifold generation are provided at https://github.com/MICA-MNI/micaopen/tree/master/structural_manifold and https://github.com/MICA-MNI/BrainSpace, and those for wiring distance calculation are provided at https://github.com/MICA-MNI/micaopen/tree/master/manifold_features.

## Acknowledgments

Dr. Bo-yong Park was funded by the National Research Foundation of Korea (NRF-2021R1F1A1052303), Institute for Information and Communications Technology Planning and Evaluation (IITP) funded by the Korea Government (MSIT) (2020-0-01389, Artificial Intelligence Convergence Research Center, Inha University; 2021-0-02068, Artificial Intelligence Innovation Hub), and Institute for Basic Science (IBS-R015-D1). Dr. Richard A. I. Bethlehem was funded by a British Academy Post-Doctoral Fellowship and the Autism Research Trust. Dr. Edward T. Bullmore was supported by a Senior Investigator award from the National Institute of Health Research (NIHR). Dr. Boris C. Bernhardt acknowledges research support from the National Science and Engineering Research Council of Canada (NSERC Discovery-1304413), CIHR (FDN-154298, PJT-174995), SickKids Foundation (NI17-039), Azrieli Center for Autism Research (ACAR-TACC), BrainCanada (Azrieli Future Leaders), and the Tier-2 Canada Research Chairs program. The Neuroscience and Psychiatry Network (NSPN) study was funded by a Wellcome Trust award to the University of Cambridge and University College London. The data were curated and analyzed using a computational facility funded by an MRC research infra-structure award (MR/M009041/1) and supported by the NIHR Cambridge Biomedical Research Centre. The views expressed are those of the authors and not necessarily those of the NHS, the NIHR or the Department of Health and Social Care.

## Author contributions

B.P. and B.C.B. designed the experiments, analyzed the data, and wrote the manuscript. C.P. and O.B. aided with the experiments. R.A.I.B. aided data curation. E.T.B. designed the NSPN MRI study. C.P., R.A.I.B., B.M., J.S., and E.T.B. reviewed and edited the manuscript. B.P. and B.C.B. are the corresponding authors of this work and have responsibility for the integrity of the data analysis.

## Supporting Information

**Fig. S1.**
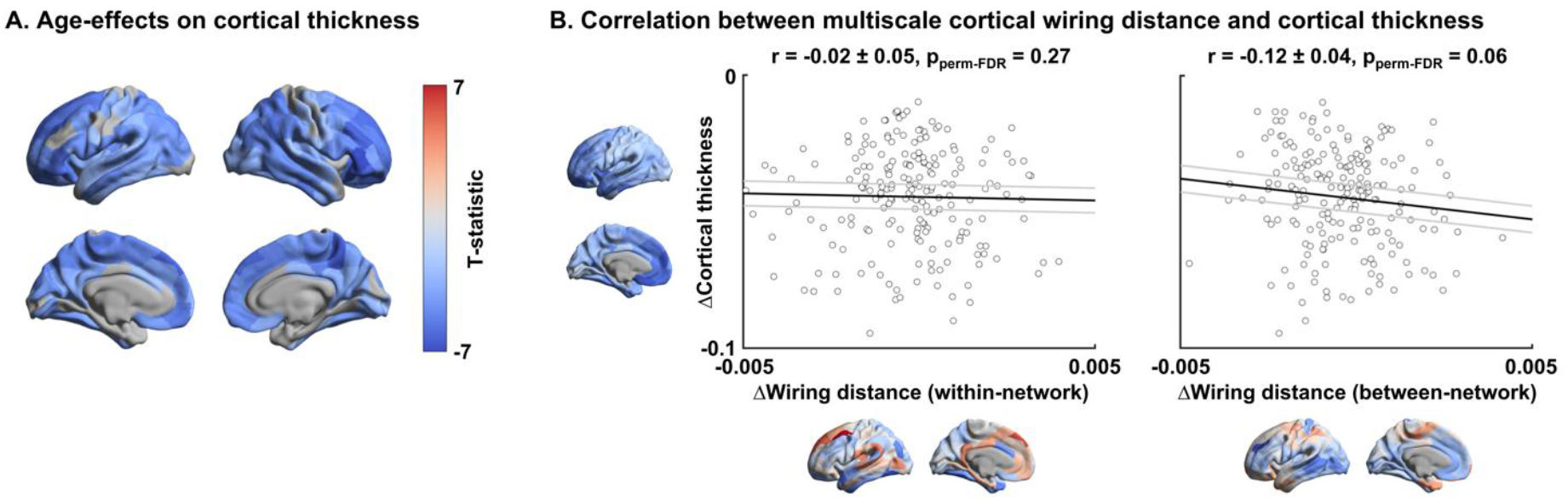
Cortical thickness effects. **(A)** The t-statistics of the identified regions that showed significant age-related changes in cortical thickness. **(B)** Linear correlations between time-related changes in cortical thickness and within/between-network wiring distance.

**Fig. S2.**
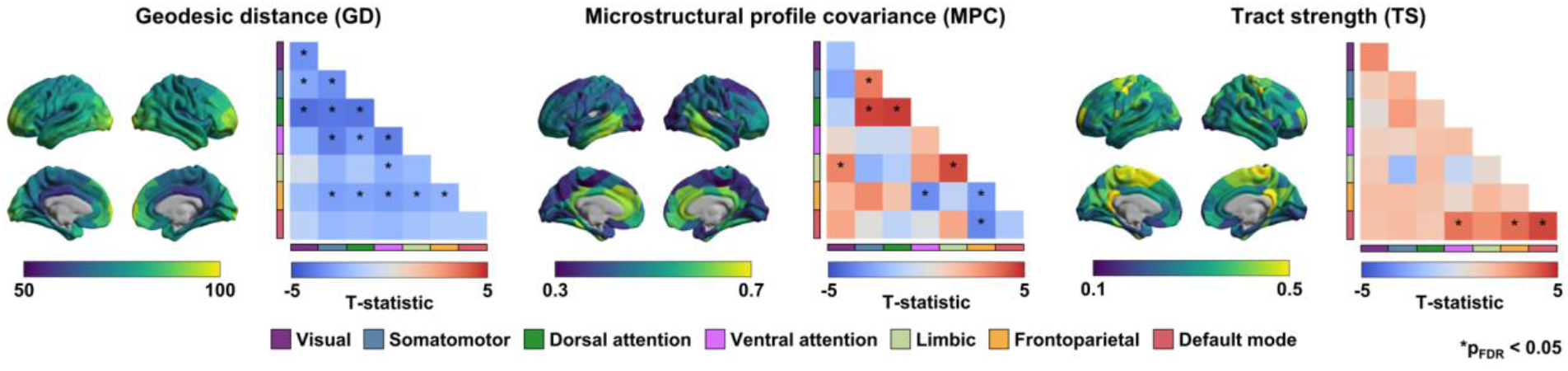
Age-effects on each cortical wiring feature. The spatial maps of GD, MPC, and TS are shown on the brain surface. The t-statistics of age-related changes on each cortical feature within- and between-networks, with significant (p_FDR_ < 0.05) results marked with asterisks. *Abbreviation:* FDR, false discovery rate.

**Fig. S3.**
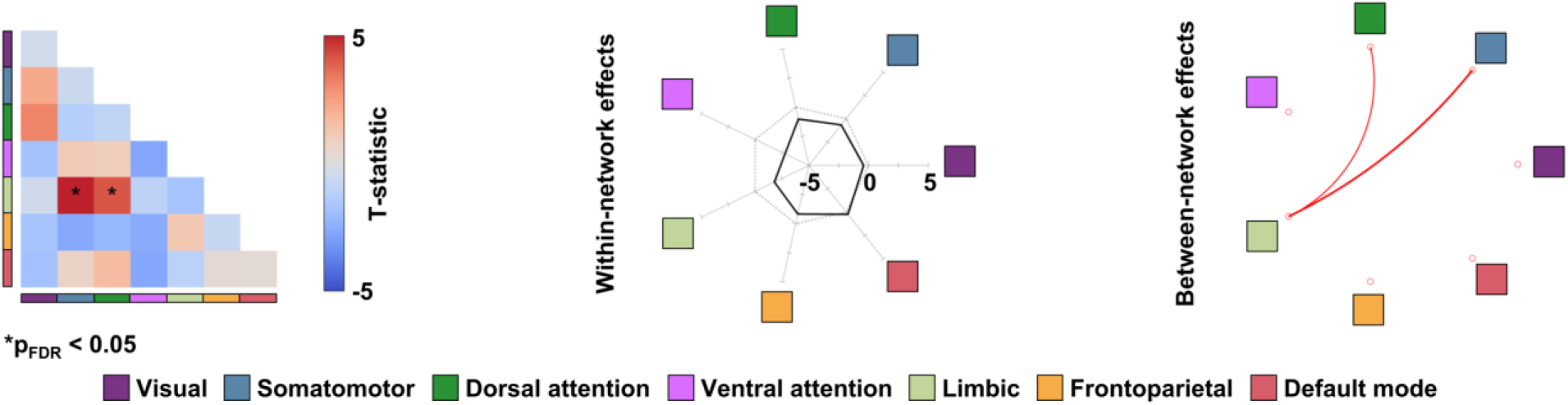
Age-effects on multiscale cortical wiring distance after controlling for mean wiring distance. The t-statistics of age-effects are reported in the matrix, and within- and between-network effects are represented with radar and circular plots, respectively. For details, see *Fig. 1*.

**Fig. S4.**
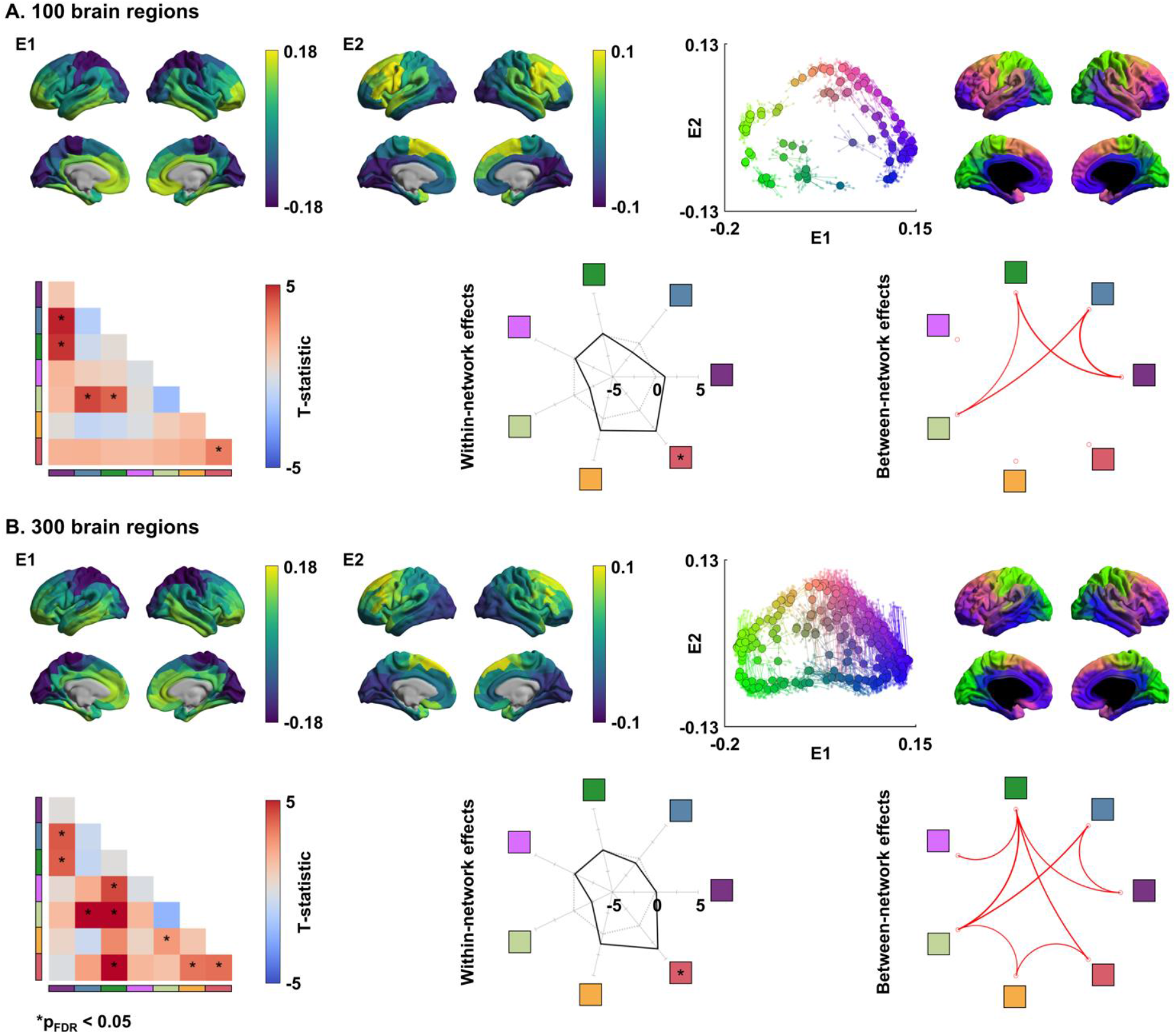
Structural manifolds and age-effects on multiscale cortical wiring distance using different parcellation scales. **(A)** Results using 100 and **(B)** 300 parcellations. Two eigenvectors (E1, E2) estimated from the cortical wiring features (top) and t-statistics of age-effects within- and between-networks (bottom) are reported. For details, see *Fig. 1*.

**Fig. S5.**
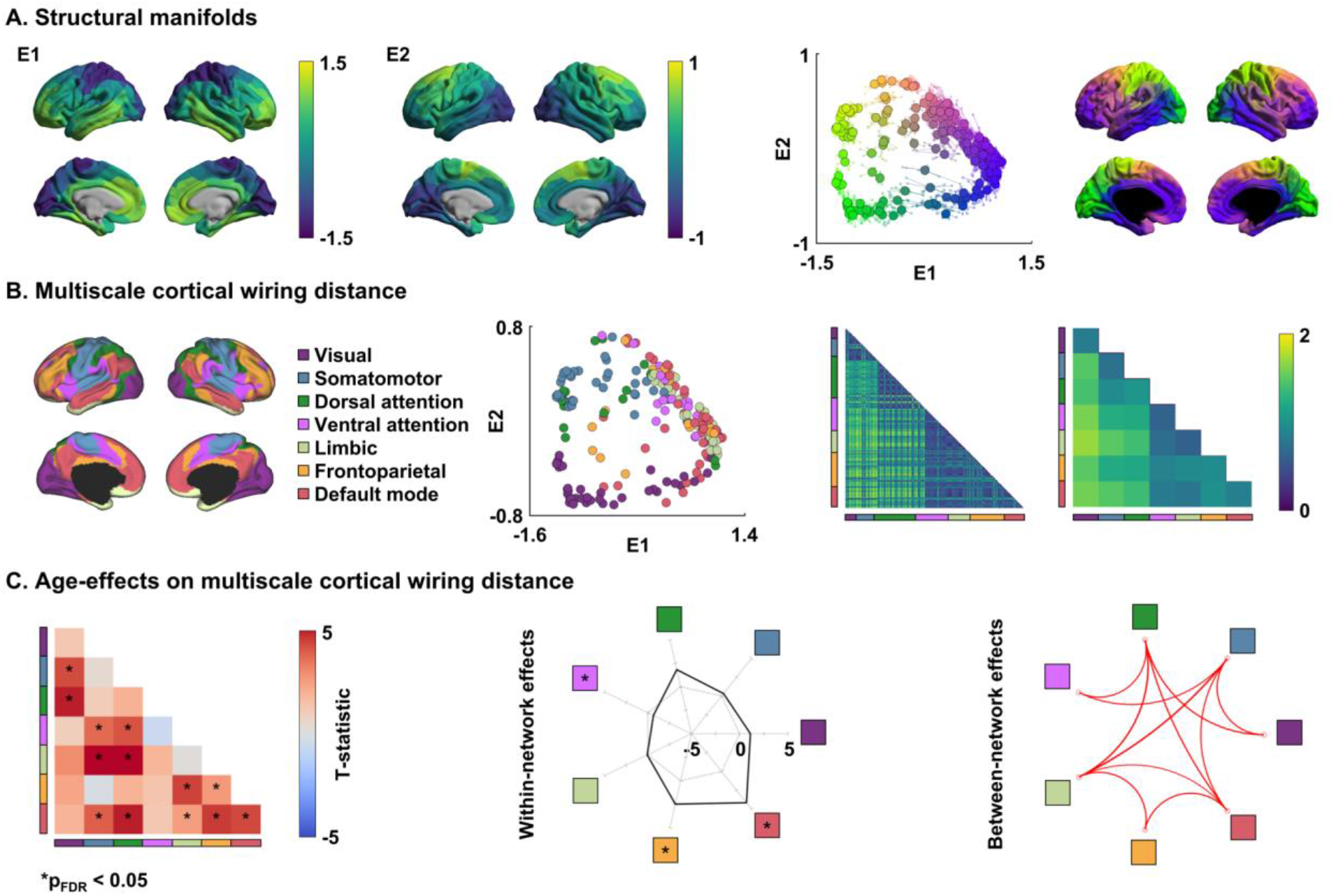
Structural manifolds derived using principal component analysis and age-effects on multiscale cortical wiring distance. **(A)** Two eigenvectors (E1, E2) estimated from the cortical wiring features. **(B)** The wiring distance summarized based on functional communities. **(C)** The t-statistics of age-effects on wiring distance within- and between-networks. For details, see *Fig. 1*.

**Fig. S6.**
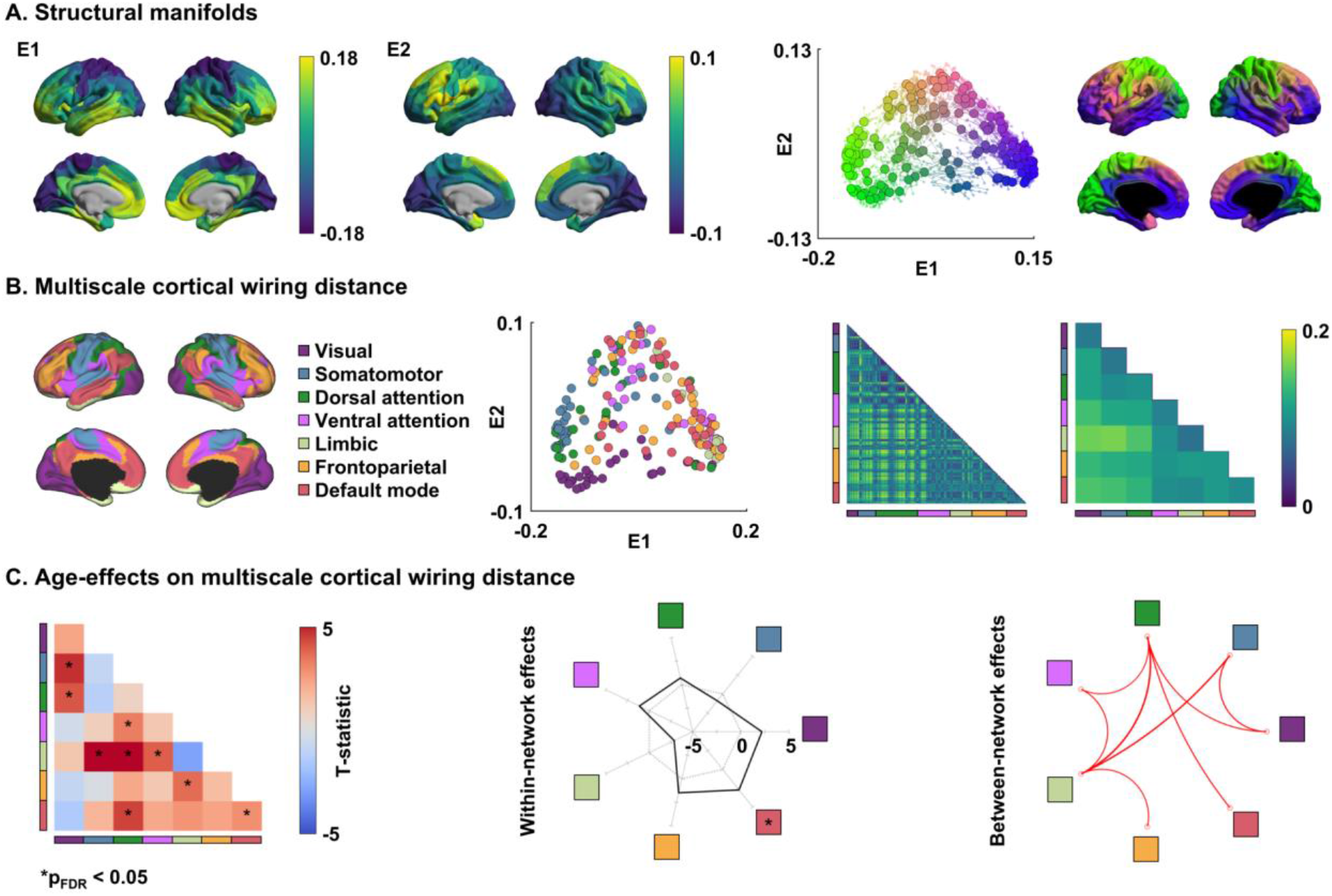
Structural manifolds and age-effects on multiscale cortical wiring distance using Schaefer 200 parcellation. **(A)** Two eigenvectors (E1, E2) estimated from the cortical wiring features. **(B)** The wiring distance summarized based on functional communities. **(C)** The t-statistics of age-effects on wiring distance within- and between-networks. For details, see *Fig. 1*.

### Neuroscience in Psychiatry Network (NSPN) Consortium author list

#### Principal investigators

Edward Bullmore (CI from 01/01/2017)^1,2,3^; Raymond Dolan^4,5^; Ian Goodyer (CI until 01/01/2017)^1^; Peter Fonagy^6^; Peter Jones^1^

#### NSPN (funded) staff

Michael Moutoussis^4,5^; Tobias Hauser^4,5^; Sharon Neufeld^1^; Rafael Romero-Garcia^1,2^; Michelle St Clair^1^; Petra Vértes^1,2^; Kirstie Whitaker^1,2^; Becky Inkster^1^; Gita Prabhu^4,5^; Cinly Ooi^1^; Umar Toseeb^1^; Barry Widmer^1^; Junaid Bhatti^1^; Laura Villis^1^; Ayesha Alrumaithi^1^; Sarah Birt^1^; Aislinn Bowler^5^; Kalia Cleridou^5^; Hina Dadabhoy^5^; Emma Davies^1^; Ashlyn Firkins^1^; Sian Granville^5^; Elizabeth Harding^5^; Alexandra Hopkins^4,5^; Daniel Isaacs^5^; Janchai King^5^; Danae Kokorikou^5,6^; Christina Maurice^1^; Cleo McIntosh^1^; Jessica Memarzia^1^; Harriet Mills^5^; Ciara O’Donnell^1^; Sara Pantaleone^5^; Jenny Scott^1^; Beatrice Kiddle^1^; Ela Polek^1^

#### Affiliated scientists

Pasco Fearon^6^; John Suckling^1^; Anne-Laura van Harmelen^1^; Rogier Kievit^4,7^; Sam Chamberlain^1^

*^1^Department of Psychiatry, University of Cambridge, United Kingdom*

*^2^Behavioural and Clinical Neuroscience Institute, University of Cambridge, United Kingdom*

*^3^ImmunoPsychiatry, GlaxoSmithKline Research and Development, United Kingdom*

*^4^Max Planck University College London Centre for Computational Psychiatry and Ageing Research, University College London, UK*

*^5^Wellcome Centre for Human Neuroimaging, University College London, United Kingdom*

*^6^Research Department of Clinical, Educational and Health Psychology, University College London, United Kingdom*

*^7^Medical Research Council Cognition and Brain Sciences Unit, University of Cambridge, United Kingdom*

